## Supplemental Figures and Tables for "SWI/SNF chromatin remodeler complex within the reward pathway is required for behavioral adaptations to stress"

Supplementary Figures and Tables

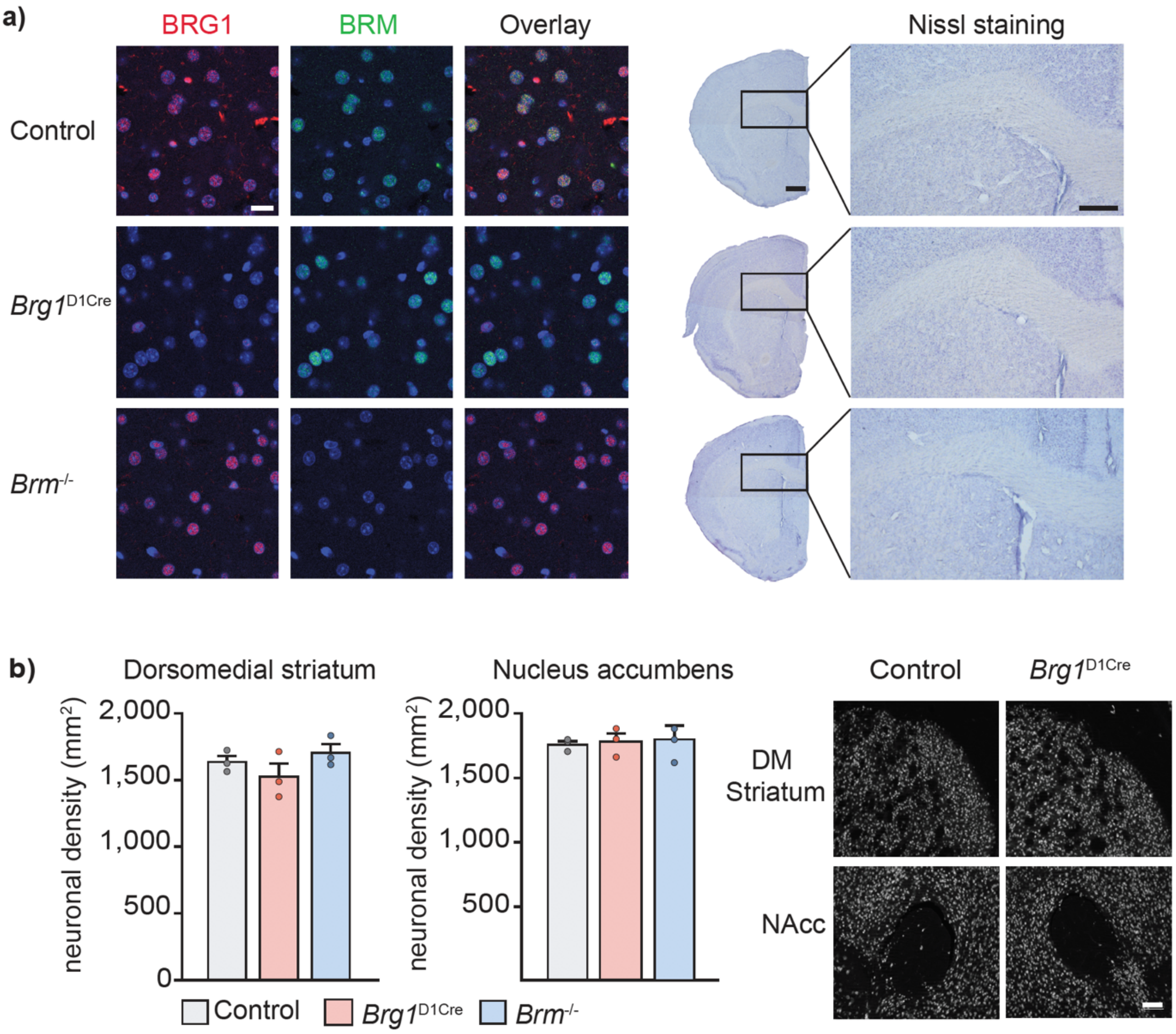


**Figure S1: Genetic inactivation of *Brg1* in dopaminoceptive neurons in *Brg1*$^{D1Cre}$ mice and of *Brm* in the whole organism in *Brm*$^{-/-}$ mice. (a)** Left panel: BRG1 (red) and BRM (green) co-staining within the nucleus accumbens of control (upper row), *Brg1*$^{D1Cre}$ (middle row), and *Brm*$^{-/-}$ (lower row) mice. DAPI nuclear staining is in blue. Scale bar = 25 µm. Right panel: Nissl staining from striatal coronal slices of control (upper row), *Brg1*$^{D1Cre}$ (middle row), and *Brm*$^{-/-}$ (lower row). Scale bar=200 µm. **(b)** Neuronal density using NeuN staining in the dorsomedial striatum and nucleus accumbens of control (n=3), *Brg1*$^{D1Cre}$ (n=3) and *Brm*$^{-/-}$ mice (n=3). Scale bar = 100 µm Data are represented as mean ± s.e.m.

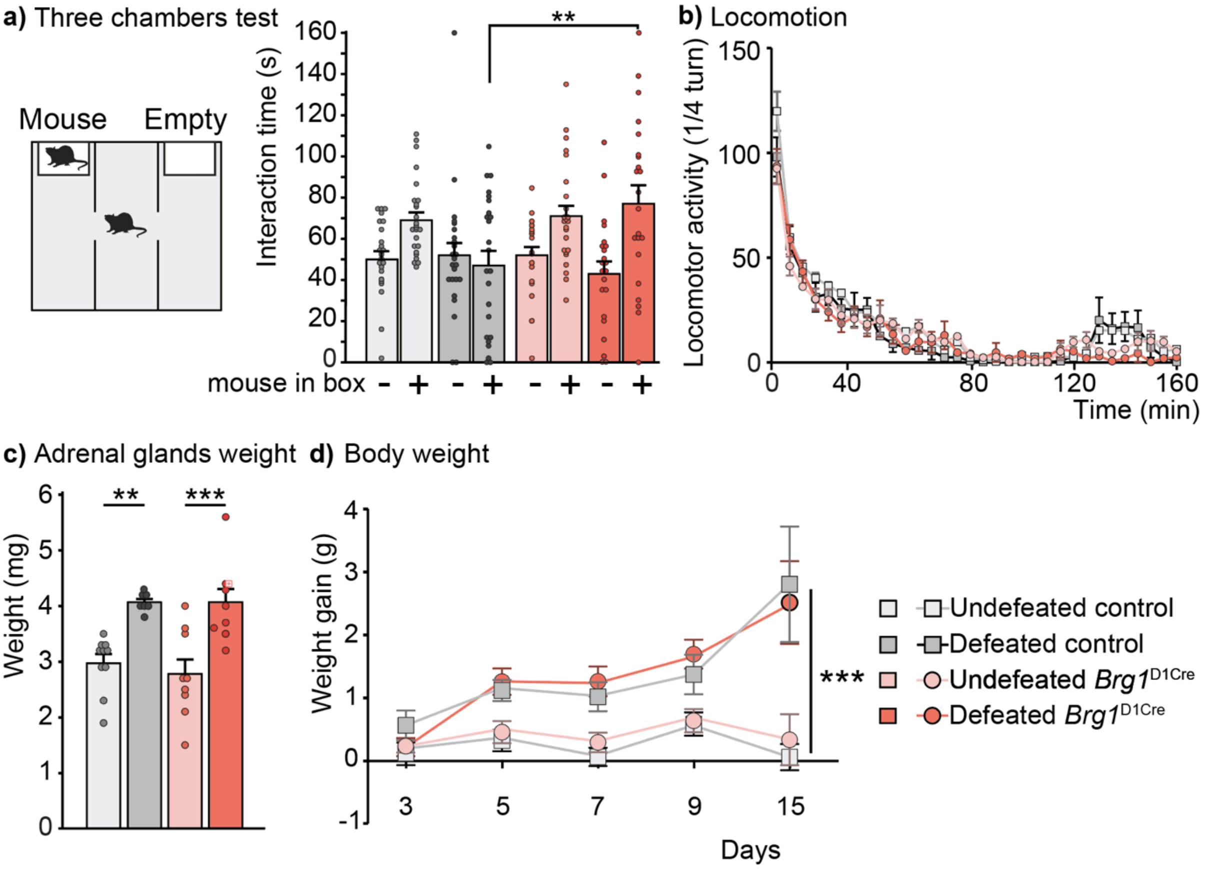


**Figure S2: Behavioral and anatomical consequences of repeated social defeat in controls and *Brg1*$^{D1Cre}$. (a)** Left panel: schematic view of the three chambers interaction test. Right panel: social interaction in the three chambers test. Undefeated control n=25, defeated control n=24, undefeated *Brg1*$^{D1Cre}$ n=24, defeated *Brg1*$^{D1Cre}$ n=22. Maine genotype effect F(1,91)=5.1; p<0.05. Main stress effect F(1,91)=4.32; p<0.05. Main social target effect F(1,91)=11.9; p<0.001. Three-way ANOVA with repeated measure followed by Bonferroni correction **p<0.01. **(b)** Effect of repeated social defeat on locomotor activity in control and *Brg1*$^{D1Cre}$ mice. Undefeated control n=9, defeated control n=6, undefeated *Brg1*$^{D1Cre}$ n=8, defeated *Brg1*$^{D1Cre}$ n=9. **(c)** Adrenal glands weight in undefeated control n=10, defeated control n=7, undefeated *Brg1*$^{D1Cre}$ n=9, defeated *Brg1*$^{D1Cre}$ n=9. Main stress effect F(1,31)=32.8; p<0.001. Two-way ANOVA followed by Bonferroni correction control undefeated vs control defeated **p<0.01; *Brg1*$^{D1Cre}$ undefeated vs *Brg1*$^{D1Cre}$ defeated ***p<0.001. **(d)** Body weight gain from the day prior repeated social defeat in undefeated control n=12, defeated control n=12, undefeated *Brg1*$^{D1Cre}$ n=13, defeated *Brg1*$^{D1Cre}$ n=13. Three-way ANOVA with repeated measure, interaction stress x days F(4,184)=10.37; ***p<0.001. Data are represented as mean ± s.e.m.

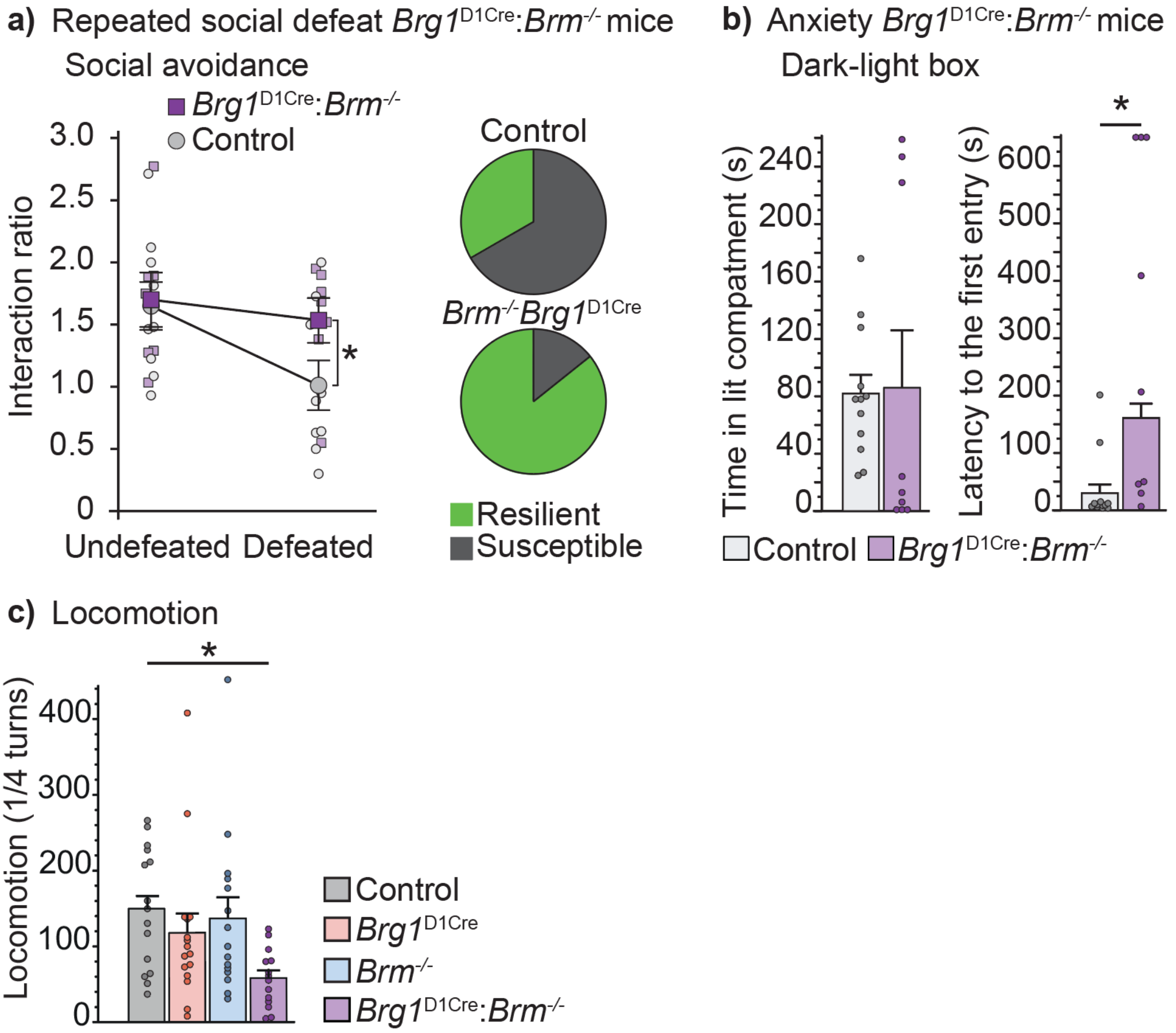


**Figure S3: Behavioral consequences of repeated social defeat in controls and compound mutants. (a)** Effect of repeated social defeat on social interaction time in control and *Brg1*$^{D1Cre}$:*Brm*$^{-/-}$ mice (left panel). Note that the same mice have been tested before and after repeated social defeat. Hence the data are represented in terms of ratio between the interaction time with a mouse and the time spent in the vicinity of the box while empty. n=9 (control mice); n=7 (compound mutant mice). Main stress effect $F(1,14)=5.093$; $p<0.05$. Two-way ANOVA with repeated measure followed by Bonferroni correction control unstressed vs control stressed $^*p<0.05$. Right panel, percentage of resilient (in green) and susceptible mice (in gray) in defeated controls (n=9) and defeated *Brg1*$^{D1Cre}$:*Brm*$^{-/-}$ mice (n=7). $X^2(1,16)=4.39$; $p<0.05$. **(b)** Anxiety-like behavior of *Brg1*$^{D1Cre}$:*Brm*$^{-/-}$ and control mice in the dark-light box test with the time in the lit compartment (left panel) and the latency of first entry in the lit compartment (right panel). Controls n=12, compound mutants n=9. Mann-

Whitney test for latency $p<0.01$. **(c)** Basal locomotor activity during 30 minutes in control (n=15), *Brg1*$^{D1Cre}$ (n=16), *Brm*$^{-/-}$ (n=15) and *Brg1*$^{D1Cre}$:*Brm*$^{-/-}$ mice (n=14). One-way ANOVA $F(3,56)=3.24$ followed by Bonferroni correction control vs *Brg1*$^{D1Cre}$:*Brm*$^{-/-}$ *$p<0.05$. Data are represented as mean ± s.e.m.

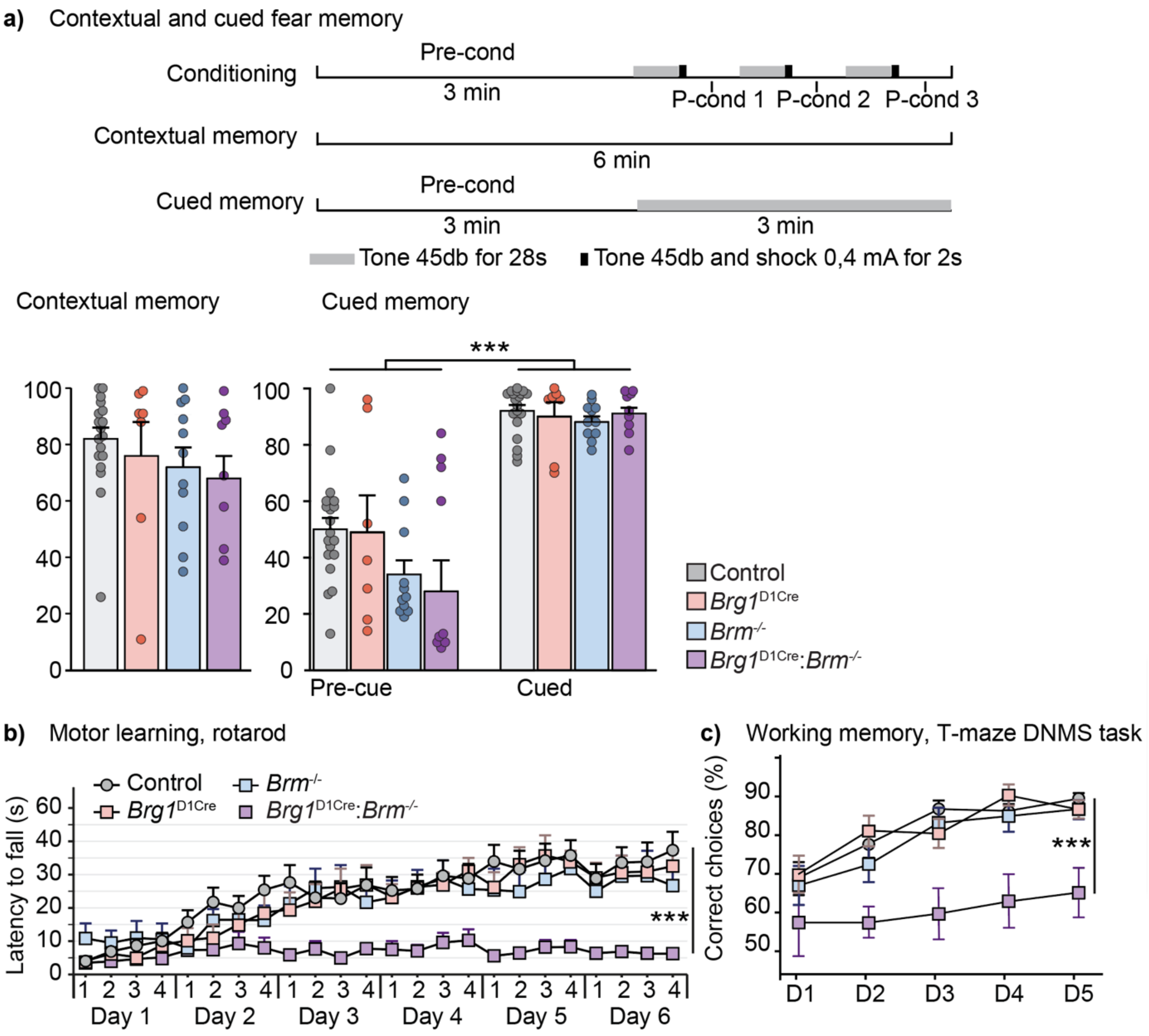


**Figure S4: Memory and motor coordination in controls, *Brg1*$^{\text{D1Cre}}$, *Brm-/-* and compound mutant mice. (a)** Fear memory in control (n=19), *Brg1*$^{\text{D1Cre}}$ mice (n=7), *Brm*$^{-/-}$ (n=11) and *Brg1*$^{\text{D1Cre}}$:*Brm*$^{-/-}$ mice (n=9). Schematic of the fear conditioning protocol with the conditioning (up), the contextual memory (middle) and the cued memory (bottom). Lower left chart, freezing percentage during the contextual fear memory test. One-way ANOVA $F(3,42)=0.83$; $p=0.48$. Lower right chart, freezing percentage during the cued fear memory test. Two-way ANOVA with repeated measure, main cue effect $F(1,42)=219.7$; $p<0.001$. **(b)** Motor coordination and learning in control (n=12), *Brg1*$^{\text{D1Cre}}$ (n=11), Brm$^{-/-}$ (n=10) and *Brg1*$^{\text{D1Cre}}$:*Brm*$^{-/-}$ mice (n=10). Interaction time x genotype $F(23,897)=2.97$; $p<0.001$. Two-way repeated ANOVA with repeated measure followed by Bonferroni correction Day 6-4 control vs compound mutant ***$p<0.001$. **(c)** T-Maze non-matching to sample working memory task in control (n=22), *Brg1*$^{\text{D1Cre}}$ mice (n=13), *Brm*$^{-/-}$ (n=11) and *Brg1*$^{\text{D1Cre}}$:*Brm*$^{-/-}$ mice (n=9). Main days effect $F(4,204)=17.3$; $p<0.001$. Main genotype effect $F(3,51)=9.91$; $p<0.001$. Two-way ANOVA with repeated measure followed by Bonferroni correction control vs compound mutant **$p<0.01$, ***$p<0.001$. Data are represented as mean ± s.e.m.

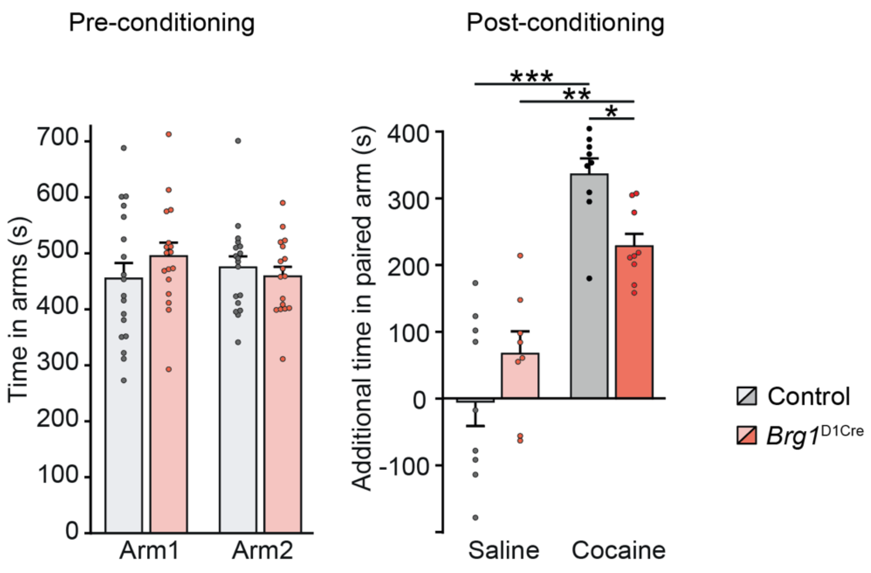


**Figure S5: Conditioned place preference to cocaine in control and *Brg1*$^{\mathbf{D1Cre}}$ mice**. (Left chart) Time spent in each compartment of the conditioned place preference setup during the preconditioning phase. Control n=18, *Brg1*$^{D1Cre}$ n=17. (Right chart) Additional time spent in the saline- and cocaine-paired compartment after conditioning in controls and *Brg1*$^{D1Cre}$ mice. Saline- and cocaine-conditioned controls n=9 for each, saline- and cocaine-conditioned *Brg1*$^{D1Cre}$ mice n=8 and n=9 respectively. Main drug effect $F(1,31)=68.8$; $p<0.001$. Genotype x drug $F(1,31)=8.683$; $p<0.01$. Two-way ANOVA followed by Bonferroni correction saline vs cocaine ***$p<0.001$, **$p<0.01$; control vs *Brg1*$^{D1Cre}$ *$p<0.05$. Data are represented as mean ± s.e.m.

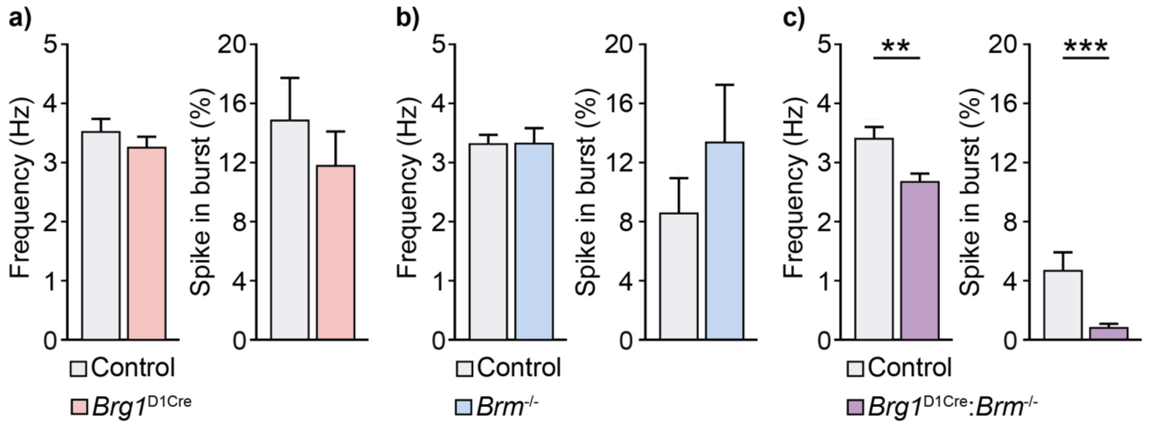


**Figure S6: Basal spontaneous activity of ventral tegmental area dopamine neurons in control, *Brg1*$^{D1Cre}$, *Brm*$^{-/-}$ and *Brg1*$^{D1Cre}$:*Brm*$^{-/-}$ mice. (a)** Firing frequency (left chart) and percentage of spike in burst (right chart) in control (n=49) and BRG1$^{D1Cre}$ mice (n=81) (data from Figure 4a and b). **(b)** Firing frequency (left chart) and percentage of spike in burst (right chart) in control (n=48) and *Brm*$^{-/-}$ mice (n=38). **(c)** Firing frequency (left chart) and percentage of spike in burst (right chart) in control (n=49) and *Brg1*$^{D1Cre}$:*Brm*$^{-/-}$ mice (n=60). T-test (for frequency) and Mann-Whitney test (spike in burst) **p<0.01, ***p<0.001. Data are represented as mean ± s.e.m.

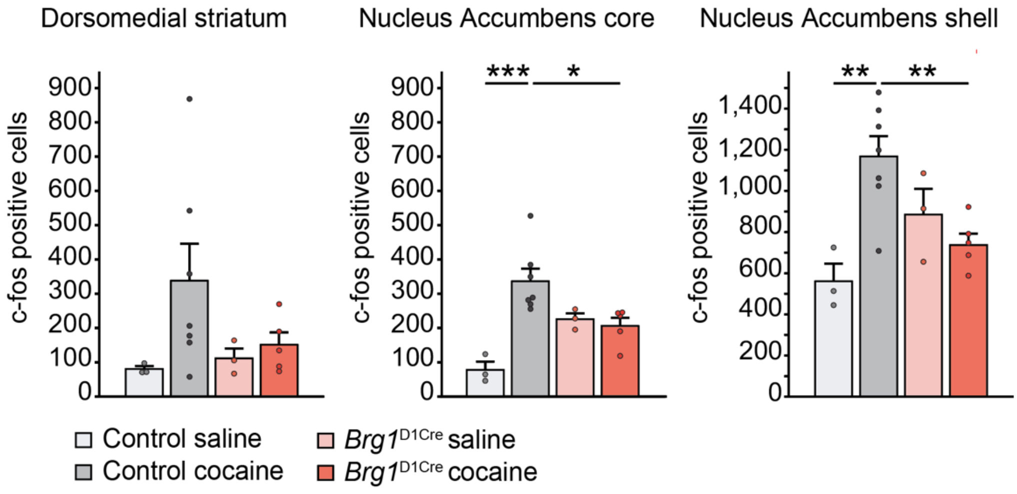


**Figure S7: *c-fos* gene induction by cocaine in controls and *Brg1*$^{D1Cre}$ mice** in the dorsomedial striatum (CPu, left chart), nucleus accumbens core (middle chart) and shell (right chart). Saline-treated control n=3, cocaine-treated control n=7, saline-treated *Brg1*$^{D1Cre}$ n=3, cocaine-treated *Brg1*$^{D1Cre}$ n=5. Two-way ANOVA followed by Bonferroni correction *p<0.05, **p<0.01. Data are represented as mean ± s.e.m.

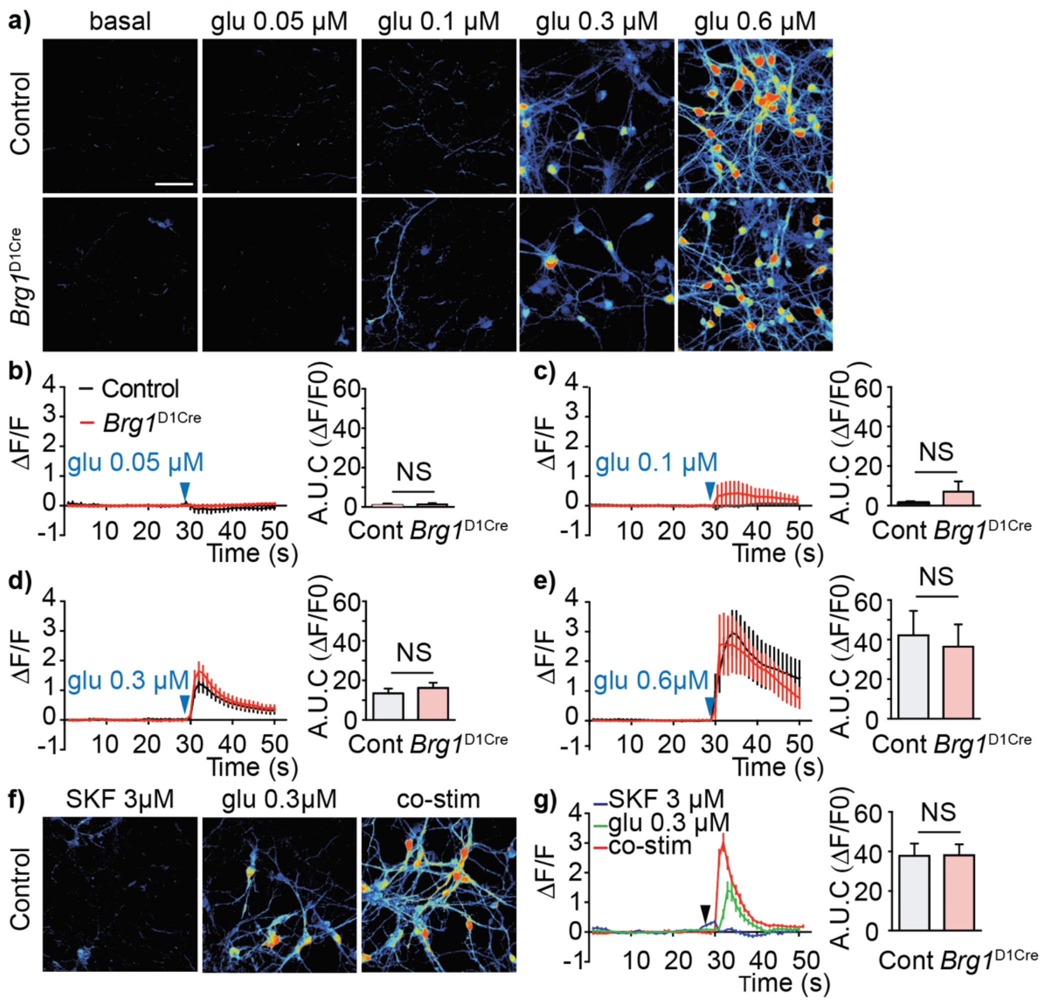


**Figure S8: Figure legend: Deletion of Brg1 in D1R-MSN preserves glutamate-mediated calcium responses as well as the integration of D1R and glutamate signals. (a)** Representative confocal images of ΔF/F calcium signals obtained at the peak of the responses during glutamate applications at the indicated doses in control (top panels) or *Brg1*$^{D1Cre}$ (bottom panels) cultured striatal neurons. Scale bar : 50 μM. **(b-e)** Left panels: dynamics of calcium signals before (0-30 sec) and during (30 to 50 sec) glutamate applications at the indicated concentrations measured from control (black) or *Brg1*$^{D1Cre}$ (red) neurons. Right panel: quantifications of the corresponding area under the curves (AUC) for control (black bars) and *Brg1*$^{D1Cre}$ (red bars) neurons. Two-way ANOVA with repeated measure, main dose effect F(4,73)=16.09 ***p<0.001; main genotype effect F(1,73)=1,01 p=0.32. **(f)** Representative pictures of the calcium responses elicited by a D1R agonist SKF38393 (3 μM) and glutamate (0.3 μM), applied separately, or together (co-stim) in control neurons. **(g)** Right panel: corresponding calcium profiles illustrating the D1R-mediated facilitation of glutamate-mediated calcium transient in control neurons. Left panel: Quantifications of the AUC of calcium transients triggered by the co-stimulation paradigm in control and *Brg1*$^{D1Cre}$. Note that the amplitude of the responses elicited by the co-stimulation is similar for both genotypes. Two-way ANOVA with repeated measure, main drug effect $F(3,96)=9.56$ ***$p<0.001$; main genotype effect $F(1,96)=1.18$ $p=0.28$; NS: not significant. Data are represented as mean ± s.e.m.

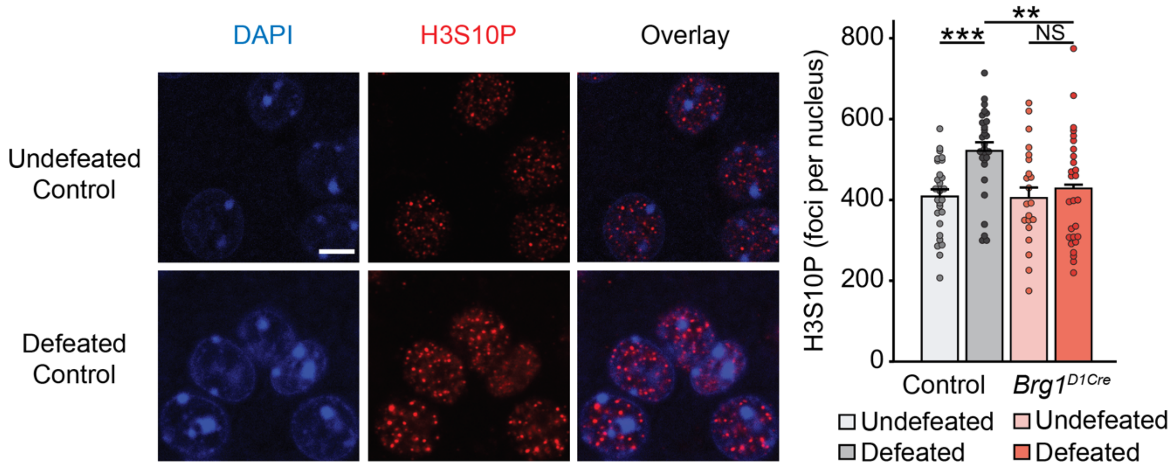


**Figure S9: Induction of H3S10P after an acute defeat in control and *Brg1*$^{\mathbf{D1Cre}}$ mice.**

Representative H3S10P staining (red) in the NAc of control mice undefeated and defeated (left panel). Scale bar = 15 µm. Induction of H3S10P by an acute social defeat in control and *Brg1*$^{D1Cre}$ mice (right panel). Undefeated control nuclei n=28 (5 mice), undefeated *Brg1*$^{D1Cre}$ nuclei = 22 (4 mice), defeated control nuclei = 28 (5 mice), defeated *Brg1*$^{D1Cre}$ nuclei = 28 (5 mice). Main genotype effect F(1,102)=4.42; p<0.05. Maine stress effect (F1,102)=8.83; p<0.01. Two-way ANOVA followed by Bonferroni correction undefeated vs defeated ***p>0.001, NS: not significant; control vs *Brg1*$^{D1Cre}$ **p<0.01. Data are represented as mean ± s.e.m.

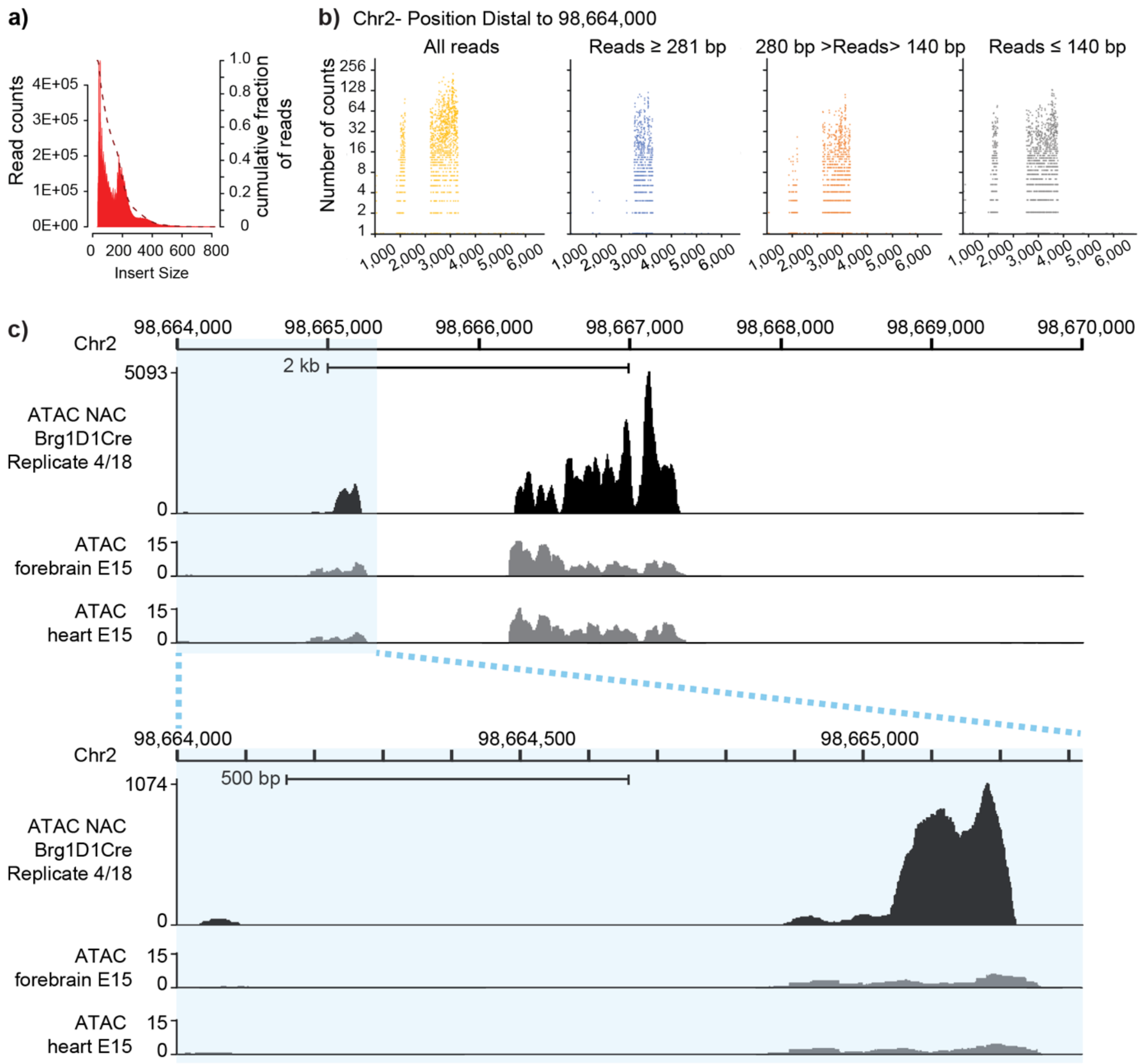


**Figure S10: Read lengths and DNA accessibility identification.** To compare data from libraries with different ratios of subnucleosomal (<=140 bp), nucleosomal (141-280 bp) and poly-nucleosomal (>=280 bp) read lengths, we first compared the qualitative and quantitative information relative to the accessibility of DNA to the Tn5 transposase by using data from the library corresponding to Mutant *Brg1*$^{D1Cre}$ Replicate 4/18 and a 6-kb region (chr2: 98,664,000-98,670,000) identified by a large read number. (a) Fragment length distributions of the library corresponding to Mutant *Brg1*$^{D1Cre}$ Replicate 4/18. The X-axis indicates read lengths, *i.e.,* the genomic distance between two Tn5 insertion sites as determined by paired-end sequencing of ATAC-seq fragments. The Y-axis indicates the density of the fraction of the reads with the indicated length. **(b)** Reads mapping within the 6-kb genomic segment (chr2: 98,664,000-98,670,000) altogether identified three regions of 70 bp, 200 bp and 1 kb as did reads obtained from P0 mouse forebrain taken as a reference profile at UCSC. **(c)** Only extremities of the subnucleosomal and nucleosomal read subsets identified the three regions. The poly-nucleosomal read subset missed the 70-bp region and identified only part of the 1 kb one. The subnucleosomal read subset displayed the more complete picture of DNA accessibility.

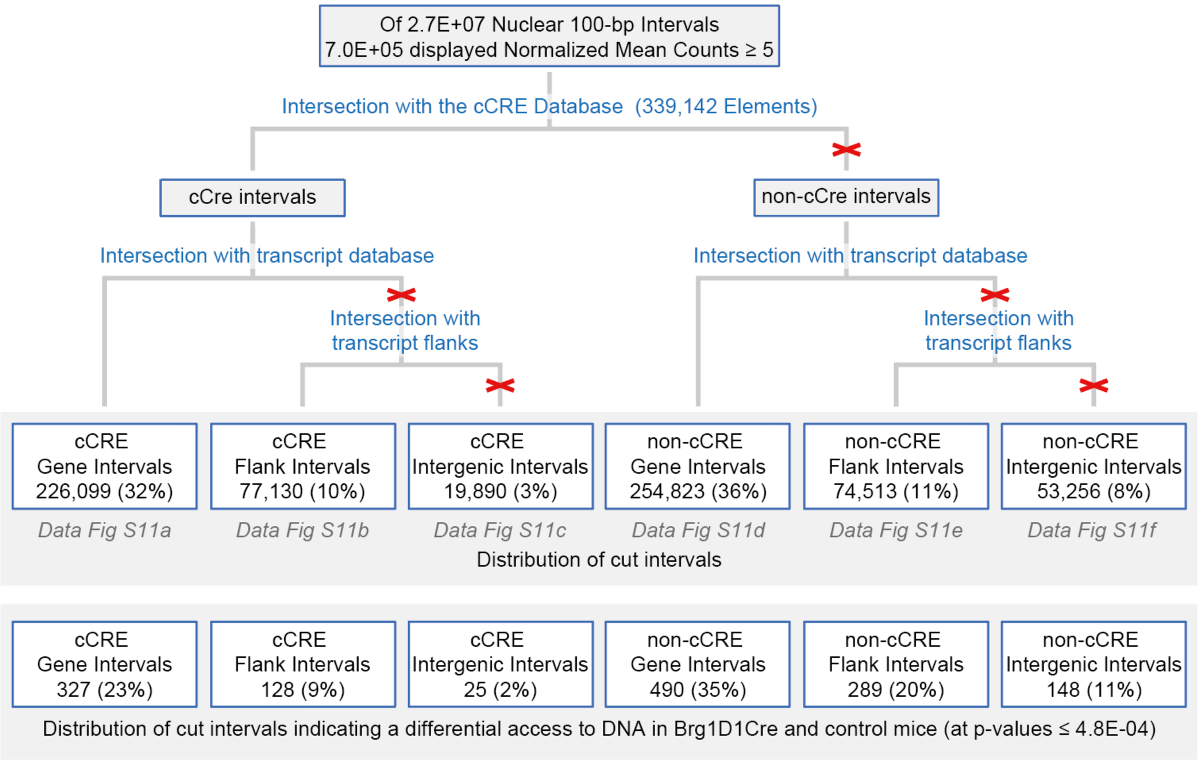


**Figure S11.** Chart of our analysis of genome accessibility. Genome accessibility was performed on 7 *Brg1*$^{D1Cre}$ and 6 control mice. The mouse genome was binned into 2.7E+7 intervals of 100 bp and the 7.1E+05 (2.4%) accessible intervals identified by a mean of 5 or more read extremities were analyzed for their overlap with cCREs, genes and gene-flanking sequences (corresponding accompanying Data files are indicated in grey letters).

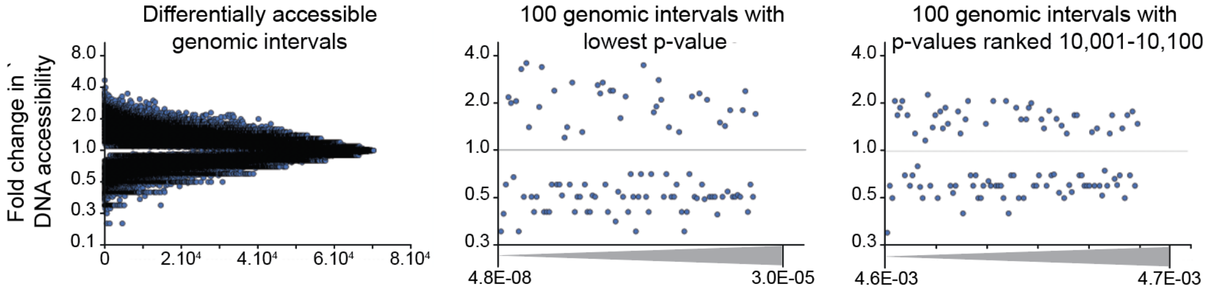


**Figure S12. Glances at fold-changes in DNA accessibility between mutant and control chromatins.** Fold-changes in accessibility are pictured for all the differentially accessible genomic intervals (left panel), the 100 ones with lowest p-values ranging from 4.8E-08 to 3.0E-05 (middle panel), and for 100 higher p-values ranging from 4.6E-03 to 4.7E-03 (right panel), see accompanying Data file.

| Antibody | Dilution | Specie | Provider | Validation reference | Specificity |
|---|---|---|---|---|---|
| BRG1 | 1/1000 | Mouse (mc) | Santa Cruz (sc-17796) | Present article | Absence of immunostaining in *Brg1* mutant neurons |
| BRM | 1/500 | Rabbit (pc) | Abcam (ab15597) | Present article | Absence of immunostaining in *Brm* mutant neurons |
| c-FOS | 1/1000 | Mouse (mc) | Santa Cruz (sc-166940) | Zhang et al. 2006 [24] | Absence of immunostaining in *c-Fos* mutant neurons |
| Cre | 1/3000 | Rabbit (pc) | House made | Kellendonk et al. 1999 [22] | Absence of immunostaining in wild type animals |
| EGR1 | 1/500 | Rabbit (mc) | Abcam (ab133695) | Liu et al. 2018 [25] | Hippocampal induction with spatial learning |
| GR | 1/2000 | Rabbit (pc) | Santa Cruz (M-20, sc-1004) | Barik et al. 2013 [7] | Absence of immunostaining in GR mutant neurons |
| H3K9me3 | 1/400 | Rabbit (pc) | Abcam (ab8898) | Sanchez et al. 2019 [30] | Colocalize with a biosensor probing for H3K9me3 in cell culture |
| H3S10P | 1/1000 | Rabbit (pc) | Millipore (06-570) | Brami-Cherrier et al. 2005 [27] | Induction by cocaine and blocked by genetic inactivation of *Msk1* gene |
| HP1 | 1/1000 | Rabbit (mc) | Abcam (ab109028) | Strom et al. 2021 [29] | Absence of immunostaining after degradation of HP1 in cell culture |
| Lamin B1 | 1/1000 | Rabbit (pc) | Abcam (ab16048) | Nmezi et al. 2019 [28] | Absence in *LaminB1* mutant mice |
| NeuN | 1/200 | Rabbit (mc) | Millipore (MABN140) | Wang et al. 2019 [23] | Co-localize with other neuronal markers (VGlut2, ChaT, GAD67) |
| pERK | 1/400 | Rabbit (mc) | Cell Signaling Technology (4370) | Contesse et al. 2021 [26] | Induced by stress and blocked by glutamate afferent silencing |

**Table S1:** List of primary antibodies. Pc : polyclona l; Mc : monoclonal

| Mouse | | Reads ≤140 bp subnucleosomal (number) | 140 < reads > 281 bp nucleosomal (number) | 281 bp ≤ reads poly-nucleosomal (number) | Total reads (number) | Reads ≤140 bp subnucleosomal (%) | 140 < reads > 281 bp nucleosomal (%) | 281 bp ≤ reads poly-nucleosomal (%) | Total reads (%) |
|---|---|---|---|---|---|---|---|---|---|
| Control | 1/1 | 3,2E+07 | 6,2E+06 | 2,0E+05 | 3,8E+07 | 0,83 | 0,16 | 0,01 | 1,00 |
| | 2/3 | 2,1E+07 | 6,1E+06 | 1,4E+05 | 2,7E+07 | 0,77 | 0,22 | 0,01 | 1,00 |
| | 3/6 | 2,5E+07 | 3,0E+07 | 2,9E+06 | 5,8E+07 | 0,43 | 0,52 | 0,05 | 1,00 |
| | 4/7 | 2,8E+07 | 5,9E+06 | 2,2E+05 | 3,4E+07 | 0,82 | 0,17 | 0,01 | 1,00 |
| | 5/22 | 9,6E+05 | 1,7E+06 | 1,1E+06 | 3,8E+06 | 0,26 | 0,45 | 0,29 | 1,00 |
| | 6/23 | 1,3E+06 | 4,1E+06 | 3,0E+06 | 8,4E+06 | 0,15 | 0,49 | 0,36 | 1,00 |
| | 7/25 | 1,6E+07 | 9,6E+06 | 1,7E+06 | 2,7E+07 | 0,59 | 0,35 | 0,06 | 1,00 |
| | 8/28 | 2,8E+07 | 6,5E+06 | 5,5E+05 | 3,5E+07 | 0,80 | 0,19 | 0,02 | 1,00 |
| Brg1D1Cre | 1/2 | 1,7E+07 | 1,5E+07 | 1,2E+06 | 3,3E+07 | 0,51 | 0,45 | 0,04 | 1,00 |
| | 2/4 | 1,9E+07 | 7,6E+05 | 6,4E+04 | 2,0E+07 | 0,96 | 0,04 | 0,00 | 1,00 |
| | 3/5 | 3,3E+07 | 1,5E+07 | 6,0E+05 | 4,9E+07 | 0,68 | 0,31 | 0,01 | 1,00 |
| | 4/18 | 3,5E+07 | 2,7E+07 | 6,8E+06 | 6,9E+07 | 0,51 | 0,39 | 0,10 | 1,00 |
| | 5/20 | 2,5E+07 | 9,3E+06 | 9,6E+05 | 3,5E+07 | 0,71 | 0,26 | 0,03 | 1,00 |
| | 6/21 | 2,2E+06 | 1,0E+07 | 2,9E+06 | 1,5E+07 | 0,15 | 0,66 | 0,19 | 1,00 |
| | 7/24 | 1,1E+06 | 3,9E+06 | 2,2E+06 | 7,2E+06 | 0,15 | 0,54 | 0,31 | 1,00 |
| | 8/26 | 1,9E+07 | 1,0E+07 | 1,5E+06 | 3,1E+07 | 0,62 | 0,33 | 0,05 | 1,00 |
| | 9/27 | 2,9E+07 | 5,6E+06 | 6,3E+05 | 3,5E+07 | 0,82 | 0,16 | 0,02 | 1,00 |

**Table S2: Read length distribution.** We constructed 17 libraries encompassing 3.8E+6 to 6.9E+7 reads. Reads were aligned to the mouse mm10 reference genome, mitochondrial and nuclear PCR duplicate reads, removed, and proper pairs (excluding those mapped to unmapped contigs), selected. Unique nuclear reads displayed different ratios of subnucleosomal (≤140 bp), nucleosomal (141-280 bp) or poly-nucleosomal (≥281 bp) lengths that were compared for their qualitative and quantitative information. Control animals 5/22 and 6/23 and *Brg1*$^{D1Cre}$ animals 6/21 and 7/24 were excluded of our analysis because of their low number (9.6E+05 to 2.2E+06 reads) of reads of subnucleosomal length when compared to the others (1.7E+07 to 3.3E+07 reads).

| | Term | Count | % | p-value | Benjamini |
|---|---|---|---|---|---|
| KEGG Pathway | Long-term depression | 9 | 1,2 | 1,2E-03 | 2,7E-01 |
| | Morphine addiction | 10 | 1,4 | 4,9E-03 | 3,1E-01 |
| | Axon guidance | 12 | 1,6 | 5,2E-03 | 3,1E-01 |
| | Hippo signaling pathway | 13 | 1,8 | 6,3E-03 | 3,1E-01 |
| | Regulation of actin cytoskeleton | 16 | 2,2 | 7,4E-03 | 3,1E-01 |
| | Pancreatic secretion | 10 | 1,4 | 7,9E-03 | 3,1E-01 |
| Tissue | Brain | 353 | 48,0 | 2,0E-09 | 3,6E-07 |
| | Brain Cortex | 50 | 6,8 | 1,8E-08 | 1,6E-06 |
| | Squamous Cell Carcinoma | 5 | 0,7 | 1,3E-05 | 8,0E-04 |
| | Fetal Brain | 24 | 3,3 | 7,8E-04 | 3,0E-02 |
| | Eyeball | 22 | 3,0 | 8,4E-04 | 3,0E-02 |
| | Cerebellum | 79 | 10,7 | 1,1E-03 | 3,2E-02 |
| | Diencephalon | 23 | 3,1 | 2,6E-03 | 6,0E-02 |
| | Embryonic tail | 25 | 3,4 | 3,2E-03 | 6,0E-02 |
| | Eye | 79 | 10,7 | 3,4E-03 | 6,0E-02 |
| | Adult Thymus | 6 | 0,8 | 3,4E-03 | 6,0E-02 |
| | Adrenal Gland | 7 | 1,0 | 6,6E-03 | 1,1E-01 |

**Table S3: Gene Ontology analyses.** The 1,273 DNA intervals of lowest p-values (4.8E-08 to 4.8E-04), corresponding to 736 genes. were analyzed for gene ontologies. The number of genes involved, and the corresponding percentage are indicated, as well as the associated p-value and adjusted p-value (Benjamini).
